# Fixed versus mixed RSA: Explaining visual representations by fixed and mixed feature sets from shallow and deep computational models

**DOI:** 10.1101/009936

**Authors:** Seyed-Mahdi Khaligh-Razavi, Linda Henriksson, Kendrick Kay, Nikolaus Kriegeskorte

## Abstract

Studies of the primate visual system have begun to test a wide range of complex computational object-vision models. Realistic models have many parameters, which in practice cannot be fitted using the limited amounts of brain-activity data typically available. Task performance optimization (e.g. using backpropagation to train neural networks) provides major constraints for fitting parameters and discovering nonlinear representational features appropriate for the task (e.g. object classification). Model representations can be compared to brain representations in terms of the representational dissimilarities they predict for an image set. This method, called representational similarity analysis (RSA), enables us to test the representational feature space as is (fixed RSA) or to fit a linear transformation that mixes the nonlinear model features so as to best explain a cortical area’s representational space (mixed RSA). Like voxel/population-receptive-field modelling, mixed RSA uses a training set (different stimuli) to fit one weight per model feature and response channel (voxels here), so as to best predict the response profile across images for each response channel. We analysed response patterns elicited by natural images, which were measured with functional magnetic resonance imaging (fMRI). We found that early visual areas were best accounted for by shallow models, such as a Gabor wavelet pyramid (GWP). The GWP model performed similarly with and without mixing, suggesting that the original features already approximated the representational space, obviating the need for mixing. However, a higher ventral-stream visual representation (lateral occipital region) was best explained by the higher layers of a deep convolutional network, and mixing of its feature set was essential for this model to explain the representation. We suspect that mixing was essential because the convolutional network had been trained to discriminate a set of 1000 categories, whose frequencies in the training set did not match their frequencies in natural experience or their behavioural importance. The latter factors might determine the representational prominence of semantic dimensions in higher-level ventral-stream areas. Our results demonstrate the benefits of testing both the specific representational hypothesis expressed by a model’s original feature space and the hypothesis space generated by linear transformations of that feature space.

**Highlights:**

1. We tested computational models of representations in ventral-stream visual areas.
2. We compared representational dissimilarities with/without linear remixing of model features.
3. Early visual areas were best explained by shallow – and higher by deep – models.
4. Unsupervised shallow models performed better without linear remixing of their features.
5. A supervised deep convolutional net performed best with linear feature remixing.

## Introduction

Sensory processing is thought to rely on a sequence of transformations of the input. At each stage, a neuronal population-code re-represents the relevant information in a format more suitable for subsequent brain computations that ultimately contribute to adaptive behavior. The challenge for computational neuroscience is to build models that perform such transformations of the input and to test these models with brain-activity data.

Here we test a wide range of candidate computational models of the representations along the ventral visual stream, which is thought to enable object recognition. The ventral stream culminates in the inferior temporal cortex, which has been intensively studied in primates (Bell et al., 2009; Hung et al., 2005; Kriegeskorte et al., 2008a) and humans (e.g. Kanwisher et al., 1997; Haxby et al., 2001; Huth et al., 2012). The representation in this higher visual area is the result of computations performed in stages across the hierarchy of the visual system. There has been good progress in understanding and modelling early visual areas (e.g. Hegdé & Van Essen, 2000; Eichhorn et al., 2009; Kay et al., 2013; Güçlü and van Gerven, 2014), and increasingly also intermediate (e.g. V4) and higher ventral-stream areas (e.g. Pasupathy and Connor, 2002; Yamins et al., 2014; Khaligh-Razavi and Kriegeskorte, 2014; Cadieu et al., 2014; Grill-Spector and Weiner, 2014; Güçlü and Gerven, 2015; Kriegeskorte, 2015; Ziemba & Freeman, 2015). Here we use data from Kay et al. (2008) to test the wide range of computational models from Khaligh-Razavi & Kriegeskorte (2014) on multiple visual areas along the ventral stream. In addition, we combine the fitting of linear models used in voxel-receptive-field modelling (Kay et al., 2008) with tests of model performance at the level of representational dissimilarities (Kriegeskorte and Kievit, 2013; Kriegeskorte et al., 2008b; Nili et al., 2014).

The geometry of a representation can be usefully characterized by a representational dissimilarity matrix (RDM) computed by comparing the patterns of brain activity elicited by a set of visual stimuli. To motivate this characterization, consider the case where dissimilarities are measured as Euclidean distances. The RDM then completely defines the representational geometry. Two representations that have the same RDM might differ in the way the units share the job of representing the stimulus space. However, the two representations would contain the same information and, down to an orthogonal linear transform, in the same format. Assuming that the noise is isotropic, a linear or radial-basis function readout mechanism could access all the same features in each of the two representations, and at the same signal-to-noise ratio.

In the framework of representational similarity analysis (RSA), representations can be compared between model layers and brain areas by computing the correlation between their RDMs (Kriegeskorte, 2009; Nili et al., 2014). Each RDM contains a representational dissimilarity for each pair of stimulus-related response patterns (Kriegeskorte and Kievit, 2013; Kriegeskorte et al., 2008b). We use the RSA framework here to compare processing stages in computational models with the stages of processing in the hierarchy of ventral visual pathway.

RSA makes it easy to test “fixed” models, that is, models that have no free parameters to be fitted. Fixed models can be obtained by optimizing parameters for task performance (Yamins et al., 2014; Krizhevsky et al., 2012; LeCun et al., 2015). This approach is essential, because realistic models of brain information processing have large numbers of parameters (reflecting the substantial domain knowledge required for feats of intelligence), and brain data are costly and limited. However, we may still want to adjust our models on the basis of brain data. For example, it might be that our model contains all the nonlinear features needed to perfectly explain a brain representation, but in the wrong proportions: with the brain devoting more neurons to some representational features than to others. Alternatively, some features might have greater gain than others in the brain representation. Both of these effects can be modelled by assigning a weight to each feature (Figure 1, upper right; Khaligh-Razavi et al., 2014; Jozwik et al., 2015). If a fixed model’s RDM does not match the brain RDM, it is important to find out whether it is just the feature weighting that is causing the mismatch.

**Figure 1.**
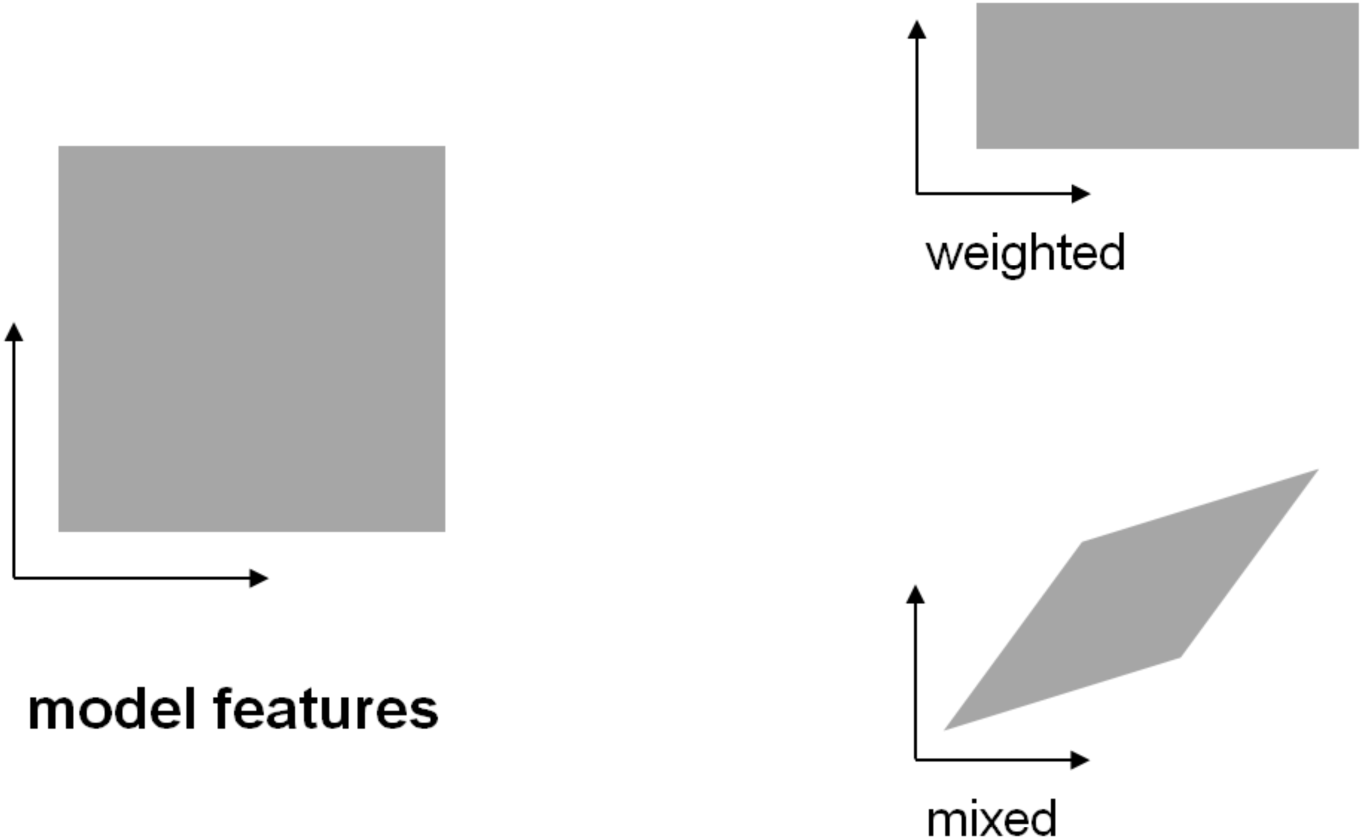
The transformation of the representational space resulting from weighting and mixing of model features. The features of a model span a representational space (left). The figure shows a cartoon 2-dimensional model-feature space. Weighting of the features (right, top) amounts to stretching and squeezing of the representational space along its original feature dimensions. Mixing (right, bottom) constitutes a more general class of transformations. We use the term mixing to denote any linear transformation, including rotation, stretching and squeezing along arbitrary axes, and shearing.

Here we take a step beyond representational weighting and explore a higher-parametric way to fit the representational space of a computational model to brain data. Our technique takes advantage of voxel/cell-population receptive-field (RF) modelling (Huth et al., 2012; Kay et al., 2008; Dumoulin and Wandell, 2008) to linearly mix model features and map them to voxel responses. Representational mixing allows arbitrary linear transformations (Figure 1, lower right). Whereas representational weighting involves fitting just one weight for each unit, representational mixing involves fitting one weight for each unit for each response channel. Where weighting can stretch or squeeze the space along its original axes, mixing can stretch and squeeze also in oblique directions, and rotate and shear the space as well. In particular, it can compute differences between the original features. Here we introduce *mixed RSA*, in which a linear remixing of the model features is first learnt using a training data set, so as to best explain the brain response patterns.

Voxel-RF modelling fits a linear transformation of the features of a computational model to predict a given voxel’s response. We bring RSA and voxel-RF modelling together by constructing RDMs based on voxel response patterns predicted by voxel-RF models. Model features are first mapped to the brain space (as in voxel-RF modelling) and the predicted and measured RDMs are then statistically compared (as in RSA). We use the linear model to predict measured response patterns for a test set of stimuli that have not been used in learning the linear remixing. We then compare the RDMs for the actual measured response patterns to RDMs for the response patterns predicted with and without linear remixing. This approach enables us to test (a) the particular representational hypothesis of each computational model and (b) the hypothesis space generated by linear transformations of the model’s computational features.

## Methods

In voxel receptive-field modelling, a linear combination of the model features is fitted using a training set and response-pattern prediction performance is assessed on a separate test set with responses to different images (Kay et al., 2008; Mitchell et al., 2008; Naselaris et al., 2011; Sprague & Serences, 2013; Cowen et al., 2014; Ester et al., 2015). This method typically requires a large training data set and also prior assumptions on the weights to prevent overfitting, especially when models have many representational features.

An alternative method is representational similarity analysis (RSA) (Kriegeskorte et al., 2008b; Kriegeskorte, 2009; Kriegeskorte and Kievit, 2013; Nili et al., 2014). RSA can relate representations from different sources (e.g. computational models and fMRI patterns) by comparing their representational dissimilarities. The representational dissimilarity matrix (RDM) is a square symmetric matrix, in which the diagonal entries reflect comparisons between identical stimuli and are 0, by definition. Each off-diagonal value indicates the dissimilarity between the activity patterns associated with two different stimuli. Intuitively, an RDM encapsulates what distinctions between stimuli are emphasized and what distinctions are de-emphasized in the representation. In this study, the fMRI response patterns evoked by the different natural images formed the basis of representational dissimilarity matrices (RDMs). The measure for dissimilarity was correlation distance (1 - Pearson linear correlation) between the response patterns. We used the RSA Toolbox (Nili et al., 2014).

The advantage of RSA is that the model representations can readily be compared with the brain data, without having to fit a linear mapping from the computational features to the measured responses. Assuming the model has no free parameters to be set using the brain-activity data, no training set of brain-activity data is needed and we do not need to worry about overfitting to the brain-activity data. However, if the set of nonlinear features computed by the model is correct, but their relative prominence or linear combination is incorrect for explaining the brain representation, classic RSA (i.e. fixed RSA) may give no indication that the model’s features can be linearly recombined to explain the representation.

Receptive-field modelling must fit many parameters in order to compare representations between brains and models. Classic RSA fits no parameters, testing fixed models without using the data to fit any aspect of the representational space. Here we combine elements of the two methods: We fit linear prediction models and then statistically compare predicted representational dissimilarity matrices.

## Mixed RSA: Combining voxel-receptive-field modelling with RSA

Using voxel-RF modelling, we first fit a linear mapping between model representations and each of the brain voxels based on a training data set (voxel responses for 1750 images) from Kay et al. (2008) (Figure 2A). We then predict the response patterns for a set of test stimuli (120 images). Finally, we use RSA to compare pattern-dissimilarities between the predicted and measured voxel responses for the 120 test images (Figure 2B). The voxel-RF fitting is a way of *mixing* the model features so as to better predict brain responses. By mixing model features we can investigate the possibility that all essential nonlinearities are present in a model, and they just need to be appropriately linearly combined to approximate the representational geometry of a given cortical area. By linear mixing of features (affine transformation of the model features), we go beyond stretching and squeezing the representational space along its original axes (Khaligh-Razavi and Kriegeskorte, 2014) and attempt to create new features as linear combinations of the original features. This affine linear recoding provides a more general transformation, which includes feature weighting as a special case.

**Figure 2.**
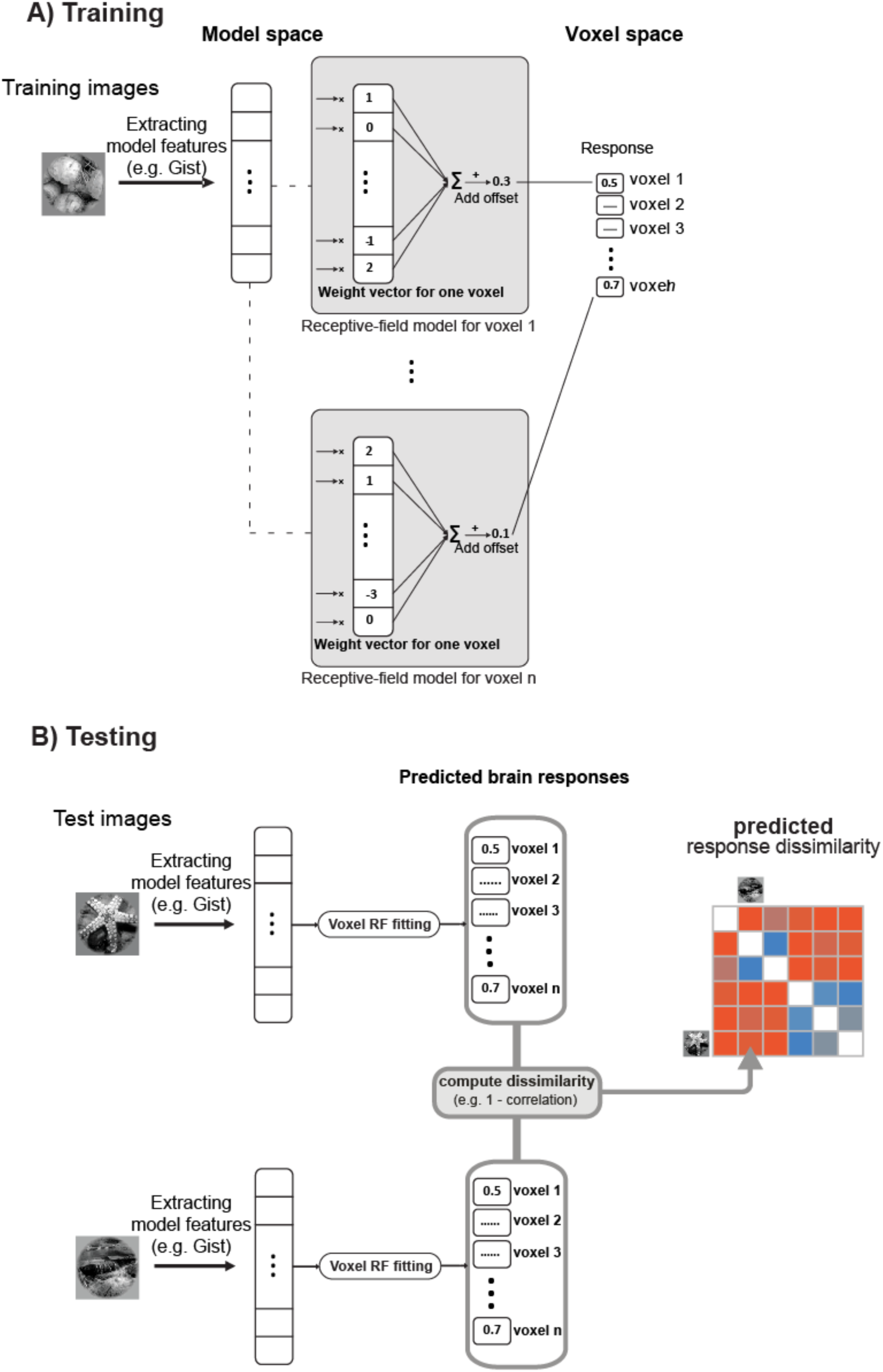
Fitting a linear model to mix representational model features. **A) Training:** learning receptive field models that map model features to brain voxel responses. There is one receptive field model for each voxel. In each receptive field model, the weight vector and the offset are learnt in a training phase using 1750 training images, for which we had model features and voxel responses. The weights are determined by gradient descent with early stopping. The figure shows the process for a sample model (e.g. gist features); the same training/testing process was done for each of the object-vision models. The offset is a constant value that is learned in the training phase and is then added to the sum of the weighted voxel responses. One offset is learnt per voxel. **B) Testing:** predicting voxel responses using model features extracted from an image. In the testing phase, we used 120 test images (not included in the training images). For each image, model features were extracted and responses for each voxel were predicted using the receptive field models learned in the training phase. Then a representational dissimilarity matrix (RDM) is constructed using the pairwise dissimilarities between predicted voxel responses to the test stimuli.

### Training

During the training phase (Figure 2A), for each of the brain voxels we learn a weight vector and an offset value that maps the internal representation of an object-vision model to the responses of brain voxels. The offset is a constant value that is learnt in the training phase and is then added to the sum of the weighted voxel responses. One offset is learnt per voxel. We only use the 1750 training images and the voxel responses to these stimuli. The weights, and the offset value are determined by gradient descent with early stopping. Early stopping is a form of regularization (Skouras et al., 1994), where the magnitude of model parameter estimates is shrunk in order to prevent overfitting. A new mapping from model features to brain voxels is learnt for each of the object-vision models.

### Regularization details

We used the regularization suggested by Skouras et al. (1994), where the shrinkage estimator of the parameters is motivated by the gradient-descent algorithm used to minimize the sum of squared errors (therefore an L2 penalty). The regularization results from early stopping of the algorithm. The algorithm stops when it encounters a series of iterations that do not improve performance on the estimation set. Stopping time is a free parameter that is set using cross-validation. An earlier stop means greater regularization. The regularization induced by early stopping in the context of gradient descent tends to keep the sizes of weights small (and tends to not break correlations between parameters). Skouras et al. (1994) show that early stopping with gradient descent is very similar to the regularization given by ridge regression, which is a L2 penalty.

### Testing

In the testing phase (Figure 2B), we use the learned mapping to predict voxel responses to the 120 test stimuli. For a given model and a presented image, we use the extracted model features and calculate the inner product of the feature vector with each of the weight vectors that were learnt in the training phase for each voxel. We then add the learnt offset value to the results of the inner product for each voxel. This gives us the predicted voxel responses to the presented image. The same procedure is repeated for all the test stimuli. Then an RDM is constructed using the pairwise dissimilarities between predicted voxel responses to the test stimuli.

### Advantages over fixed RSA and voxel receptive-field modelling

Considering the predictive performance of either (a) the particular set of features of a model (fixed RSA) or (b) linear transformations of the model (voxel-RF modeling) provides ambiguous results. In fixed RSA (Figure 3A), it remains unclear to what extent fitting a linear transformation might improve performance. In voxel-RF modelling, it remains unclear whether the set of features of the model, as is, already span the correct representational space. Mixed RSA (Figure 3B) enables us to compare fitted and unfitted variants of each model.

**Figure 3.**
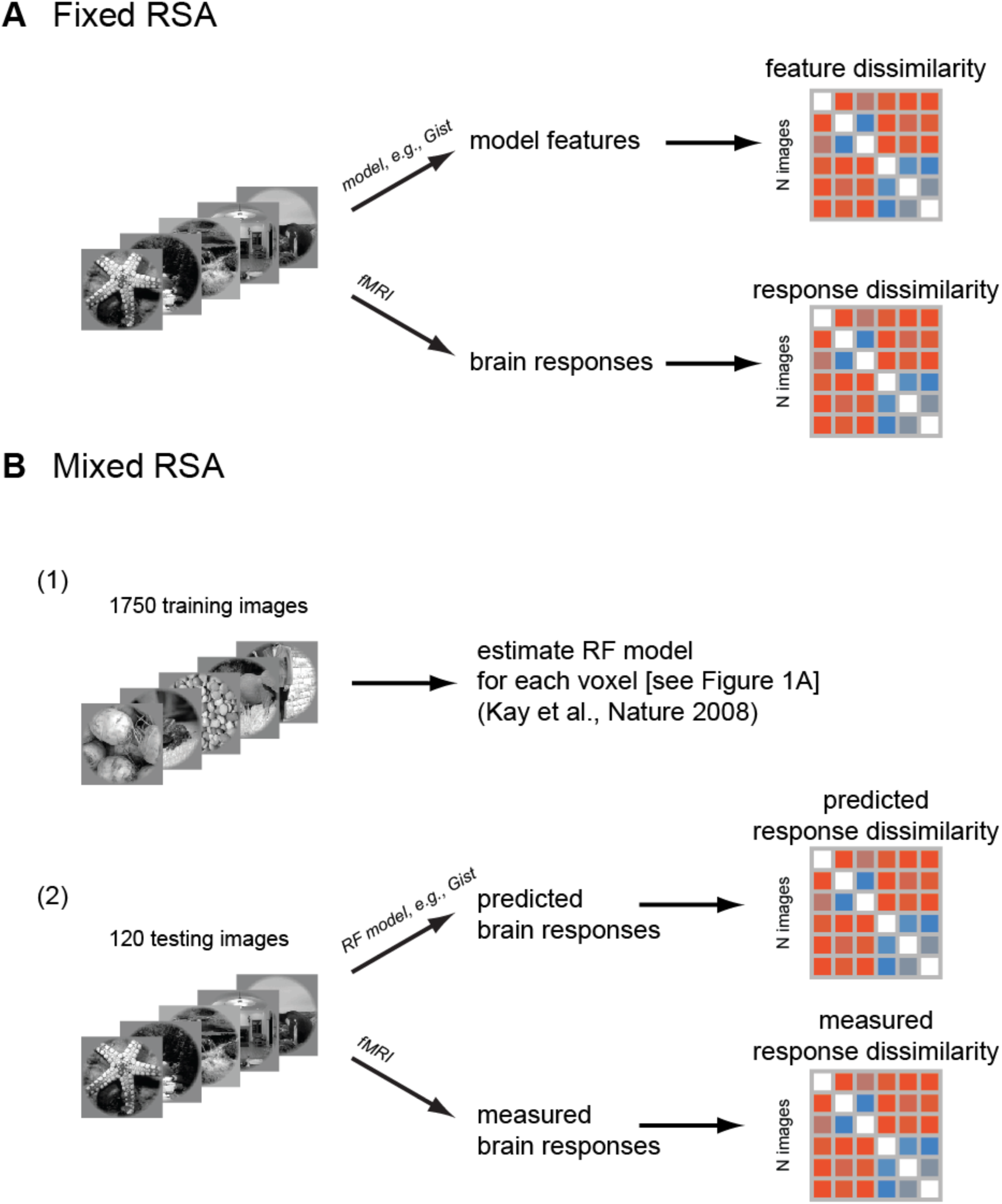
Fixed versus mixed RSA. **A) Fixed RSA**: a brain RDM is compared to a model RDM which is constructed from the model features. The model features are extracted from the images and the RDM is constructed using the pairwise dissimilarities between the model features. **B) Mixed RSA:** a brain RDM is compared to a model RDM which is constructed from mixed model features obtained via receptive field modelling (see also Figure 2). There is first a training phase in which the receptive field models are estimated for each voxel (similar to Kay et al. 2008). In the testing phase, using the learned receptive field models, voxel responses to new stimuli are predicted. Then the RDM of the predicted voxel responses is compared with the RDM of the actual measured brain (voxel) responses.

Fitted linear feature combinations may not explain the brain data in voxel-RF modelling for a combination of three reasons: (1) the features do not provide a sufficient basis, (2) the linear model suffers from overfitting, (3) the prior implicit to the regularisation procedure prevents finding predictive parameters. Comparing fitted and unfitted models in terms of their prediction of dissimilarities provides additional evidence for interpreting the results. When the unfitted model outperforms the fitted model, this suggests that the original feature space provides a better estimate of relative prominence and linear mixing of the features than the fitting procedure can provide (at least given the amount of training data used).

The method of mixed RSA, which we use here, compares representations between models and brain areas at the level of representational dissimilarities. This enables direct testing of unfitted models and straightforward comparisons between fitted and unfitted models. The same conceptual question could be addressed in the framework of voxel-RF modelling. This would require fitting linear models to the voxels with the constraint that the resulting representational space spanned by the predicted voxel responses reproduce the representational dissimilarities of the model’s original feature space as closely as possible.

## Stimuli, response measurements, and RDM computation

In this study we used the experimental stimuli and fMRI data from Kay et al. (2008); also used in Naselaris et al., (2009); Kay et al., (2011); Guclu & van Gerven (2015). The stimuli were gray-scale natural images. The training stimuli were presented to subjects in 5 scanning sessions with 5 runs in each session (overall 25 experimental runs). Each run consisted of 70 distinct images presented two times each. The testing stimuli were 120 gray-scale natural images. The data for testing stimuli were collected in 2 scanning sessions with 5 runs in each session (overall 10 experimental runs). Each run consisted of 12 distinct images presented 13 times each.

We had early visual areas (i.e. V1, V2), intermediate level visual areas (V3, V4), and LO as one of the higher visual areas. The RDMs for each ROI were calculated based on 120 test stimuli presented to the subjects. For more information about the data set, and images see supplementary methods or refer to (Kay et al., 2008; Henriksson et al., 2015).

The RDM correlation between brains ROIs and models is computed based on the 120 testing stimuli. For each brain ROI, we had ten 12×12 RDMs, one for each experimental run (10 runs with 12 different images in each = 120 distinct images overall). Each test image was presented 13 times per run. To calculate the correlation between model and brain RDMs, within each experimental run, all trials were averaged, yielding one 12×12 RDM for each run. The reported model-to-brain RDM correlations are the average RDM correlations for the ten sets of 12 images.

To judge the ability of a model RDM to explain a brain RDM, we used Kendall’s rank correlation coefficient *τ*_A_ (which is the proportion of pairs of values that are consistently ordered in both variables). When comparing models that predict tied ranks (e.g. category model RDMs) to models that make more detailed predictions (e.g. brain RDMs, object-vision model RDMs) Kendall’s *τ*_A_ correlation is recommended (Nili et al., 2014), because the Pearson and Spearman correlation coefficients have a tendency to prefer a simplified model that predicts tied ranks for similar dissimilarities over the true model.

### Inter-subject brain RDM correlations

The inter-subject brain RDM correlation is computed for each ROI for comparison with model-to-brain RDM correlations. This measure is defined as the average Kendall *τ*_A_ correlation of the ten 12×12 RDMs (120 test stimuli) between the two subjects (Figures 4–7).

**Figure 4.**
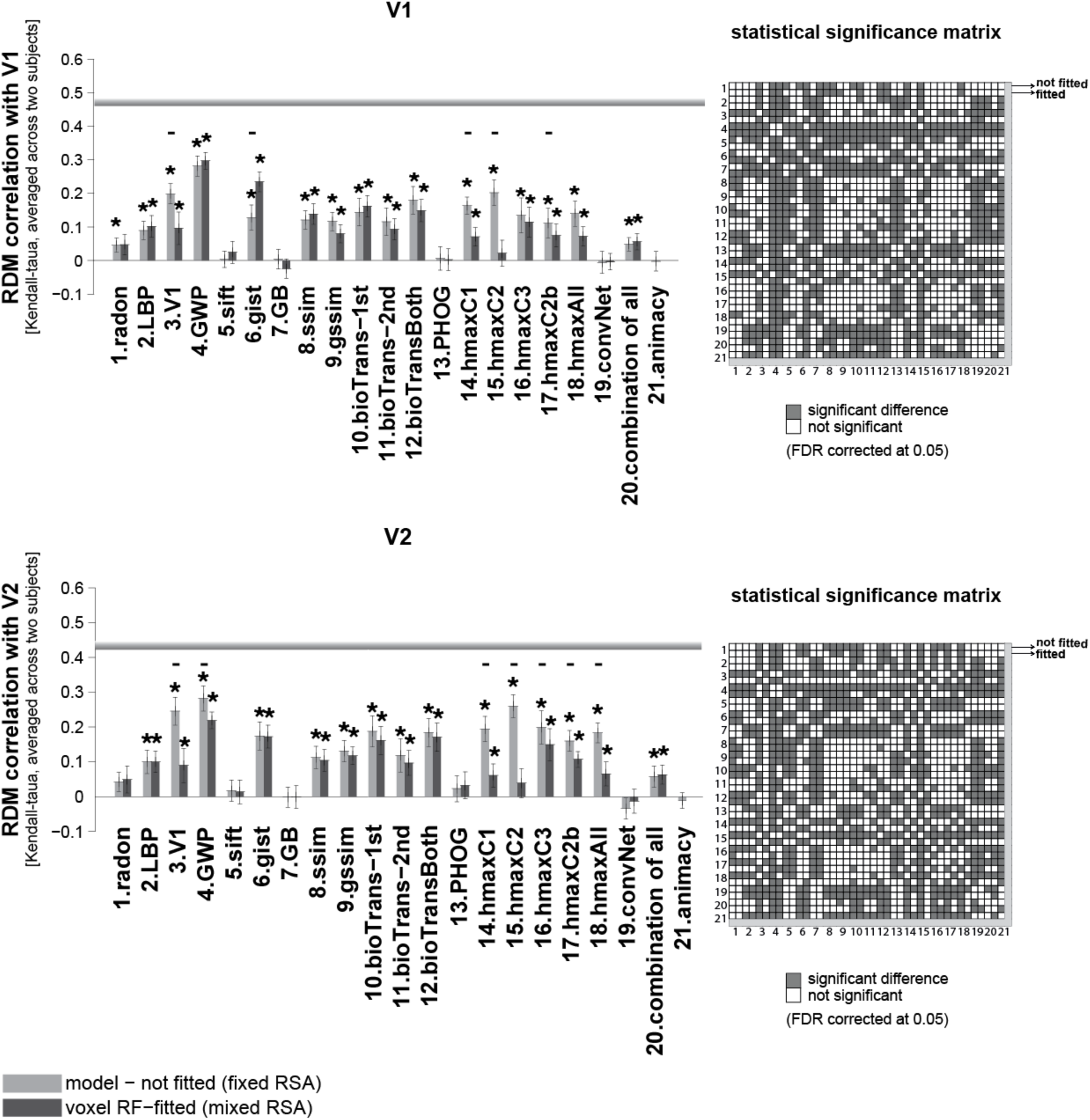
RDM correlation of unsupervised models and animacy model with early visual areas. Bars show the average of ten 12×12 RDM correlations (120 test stimuli in total) with V1, and V2 brain RDMs. There are two bars for each model. The first bar, ‘model (not fitted)’, shows the RDM correlation of a model with a brain ROI without fitting the model responses to brain voxels (fixed RSA). The second bar (voxel RF-fitted) shows the RDM correlation of a model that is fitted to the voxels of the reference brain ROI using 1750 training images (mixed RSA; refer to Figures 1 and 2 to see how the fitting is done). Stars above each bar show statistical significance obtained by signrank test (FDR corrected at 0.05). Small black horizontal bars show that the difference between the bars for a model is statistically significant (signrank test, 5% significance level - not corrected for mutiple comparisons. For FDR corrected comparison see the statistical significance matrices on the right). The results are the average over the two subjects. The gray horizontal line for each ROI indicates the inter-subject brain RDM correlation. This is defined as the average Kendall-taua correlation of the ten 12×12 RDMs (120 test stimuli) between the two subjects. The animacy model is categorical, consisting of a single binary variable, therefore mixing has no effect on the predicted RDM rank order. We therefore only show the unfitted animacy model. The color-coded statistical signficance matrices at the right side of the bar graphs show whether any of the two models perform significantly differently in explaining the corresponding reference brain ROI (FDR corrected at 0.05). Models are shown by their corresponding number; there are two rows/columns for each model, the first one representes the not-fitted version and the second one the voxel RF-fitted. A gray square in the matrix shows that the corresponding models perform signficantly differetnly in expalining the refernce brain ROI (one of them signficantly expalins the reference brain ROI better/worse).

## Models

We tested a total of 20 unsupervised computational model representations, as well as different layers of a pre-trained deep supervised convolutional neuronal network (Krizhevsky et al., 2012). In this context, by unsupervised, we mean object-vision models that had no training phase (e.g. feature extractors, such as gist), as well as models that are trained but without using image labels (e.g. HMAX model trained with some natural images). Some of the models mimic the structure of the ventral visual pathway (e.g. V1 model, HMAX); others are more broadly biologically motivated (e.g. BioTransform, convolutional networks); and the others are well-known computer-vision models (e.g. GIST, SIFT, PHOG, self-similarity features, geometric blur). Some of the models use features constructed by engineers without training with natural images (e.g. GIST, SIFT, PHOG). Others were trained in an unsupervised (e.g. HMAX) or supervised (deep CNN) fashion.

In the following sections we first compare the representational geometry of several unsupervised models with that of early to intermediate and higher visual areas using both fixed RSA and mixed RSA. We will then test a deep supervised convolutional network in terms of its ability in explaining the hierarchy of vision.

Further methodological details are explained in the supplementary materials (Supplementary methods).

## Results

### Early visual areas explained by Gabor wavelet pyramid

The Gabor wavelet pyramid (GWP) model was used in Kay et al. (2008) to predict responses of voxels in early visual areas in humans. Gabor wavelets are directly related to Gabor filters, since they can be designed for different scales and rotations. The aim of GWP has been to model early stages of visual information processing, and it has been shown that 2D Gabor filters can provide a good fit to the receptive field weight functions found in simple cells of cat striate cortex (Jones and Palmer, 1987).

The GWP model had the highest RDM correlation with both V1, and V2 (Figure 4). As for V1, the GWP model (voxel-RF fitted) performs significantly better than all other models in explaining V1 (see the ‘statistical significance matrix’). Similarly, in V2, the GWP model (unfitted) performs well in explaining this ROI. Although GWP has the highest correlation with V2, the correlation is not significantly higher than that of the V1 model, HMAX-C2, and HMAX-C3 (the ‘statistical significance matrices’ in Figure 4 show pairwise statistical comparisons between all models. The statistical comparisons are based on two-sided signed-rank test, FDR corrected at 0.05).

The GWP model comes very close to the inter-subject RDM correlation of these two early visual areas (V1, and V2), although it does not reach it. Indeed, the inter-subject RDM correlation for these two areas (V1 and V2) is much higher than those calculated for the other areas (see the inter-subject RDM correlation for V3, V4, and LO in Figures 5, 6). The highest correlation obtained between a model and a brain ROI is for the GWP model and the early visual areas V1 and V2. This suggests that early vision is better modelled or better understood, compared to other brain ROIs. It is possible that the newer Gabor-based models of early visual areas (Kay et al., 2013) explain early visual areas even better.

**Figure 5.**
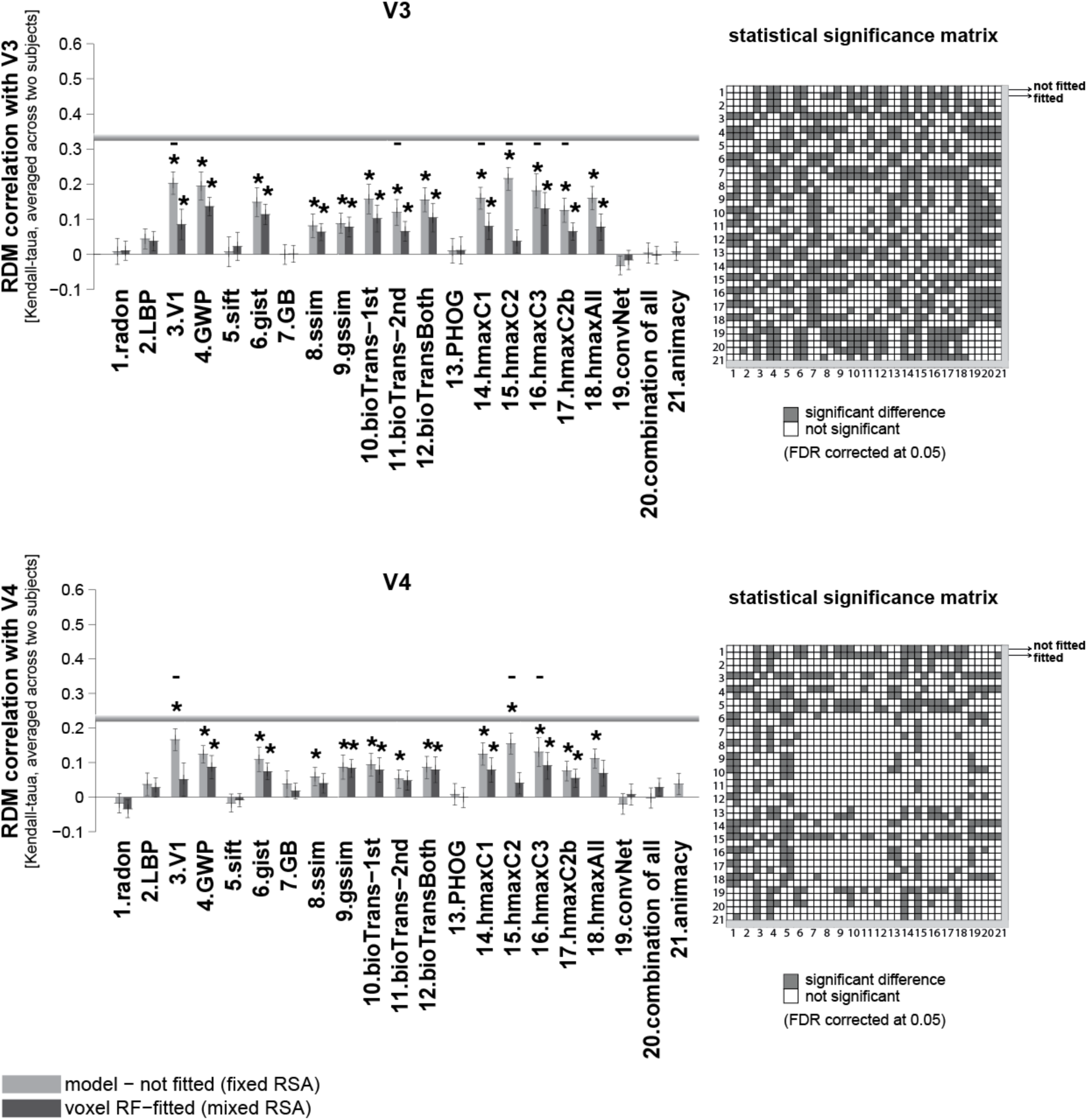
RDM correlation of unsupervised models and animacy model with intermediate-level visual areas. Bars show the average of ten 12×12 RDM correlations (120 test stimuli in total) with V3, and V4 brain RDMs. There are two bars for each model. The first bar, ‘model (not fitted)’, shows the RDM correlation of a model with a brain ROI without fitting the model responses to brain voxels (fixed RSA). The second bar (voxel RF-fitted) shows the RDM correlations of a model that is fitted to the voxels of the reference brain ROI, using 1750 training images (mixed RSA; refer to Figures 1 and 2 to see how the fitting is done). The gray horizontal line in each panel indicates inter-subject RDM correlation for that ROI. The color-coded statistical signficance matrices at the right side of the bar graphs show whether any of the two models perform significantly differently in explaining the corresponding reference brain ROI (FDR corrected at 0.05). The statistical analyses and conventions here are analogous to Figure 4.

The next best model in explaining the early visual area V1 was the voxel RF-fitted gist model. For V2, in addition to the GWP, the HMAX-C2 and C3 features also showed a high RDM correlation. Overall results suggest that shallow models are good in explaining early visual areas. Interestingly, all the mentioned models that better explained V1 and V2 are built based on Gabor-like features.

### Visual areas V3 and V4 explained by unsupervised models

Several models show high correlations with V3, and V4, and some of them come close to the inter-subject RDM correlation for V4. However, note that the inter-subject RDM correlation is lower in V4 compared to V1, V2, and V3 (Figure 5).

Intermediate layers of the HMAX model (e.g. C2 – model #15 in Figure 5) seem to perform slightly better than other models in explaining intermediate visual areas (Figure 5) –significantly better than most of the other unsupervised models (see the ‘statistical significance matrices’ in Figure 5; row/column #15 refers to the statistical comparison of HMAX-C2 features with other models—two-sided signed-rank test, FDR corrected at 0.05). More specifically, for V3, in addition to the HMAX C1, C2 and C3 features, GWP, V1 model (which is a combination of simple and complex cells), gist, and bio-transform also perform similarly well (not significantly different from HMAX-C2).

In V4, the voxel responses seem noisier (the inter-subject RDM correlation is lower); and the RDM correlation of models with this brain ROI is generally lower. The HMAX-C2 is still among the best models that explain V4 significantly better than most of the other unsupervised models. The following models perform similarly well (not significantly different from HMAX features) in explaining V4: GWP, gist, V1 model, bio-transform, and gssim (for pairwise statistical comparison between models, see the statistical significance matrix for V4).

Overall from these results we may conclude that the Gabor-based models (e.g. GWP, gist, V1 model, and HMAX) provide a good basis for predicting voxel responses in the brain from early visual areas to intermediate levels. More generally, intermediate visual areas are best accounted for by the unfitted versions of the unsupervised models. It seems that for most of the models the mixing does not improve the RDM correlation of unsupervised model features with early and intermediate visual areas.

### Higher visual areas explained by mixed deep supervised neural net layers

For the higher visual area LO (Grill-Spector et al., 2011; Mack et al., 2013), a few of the unsupervised models explained a significant amount of non-noise variance (Figure 6). These were GWP, gist, geometric blur (GB), ssim, and bio-transform (1^st^ stage). None of these models reached the inter-subject RDM correlation for LO. Animacy model achieved the highest RDM correlation (though not significantly higher than some of the other unsupervised models). The animacy model is a simple model RDM that shows the animate-inanimate distinction (it is not an image-computable model). The animacy came close to the inter-subject RDM correlation for LO, but did not reach it.

**Figure 6.**
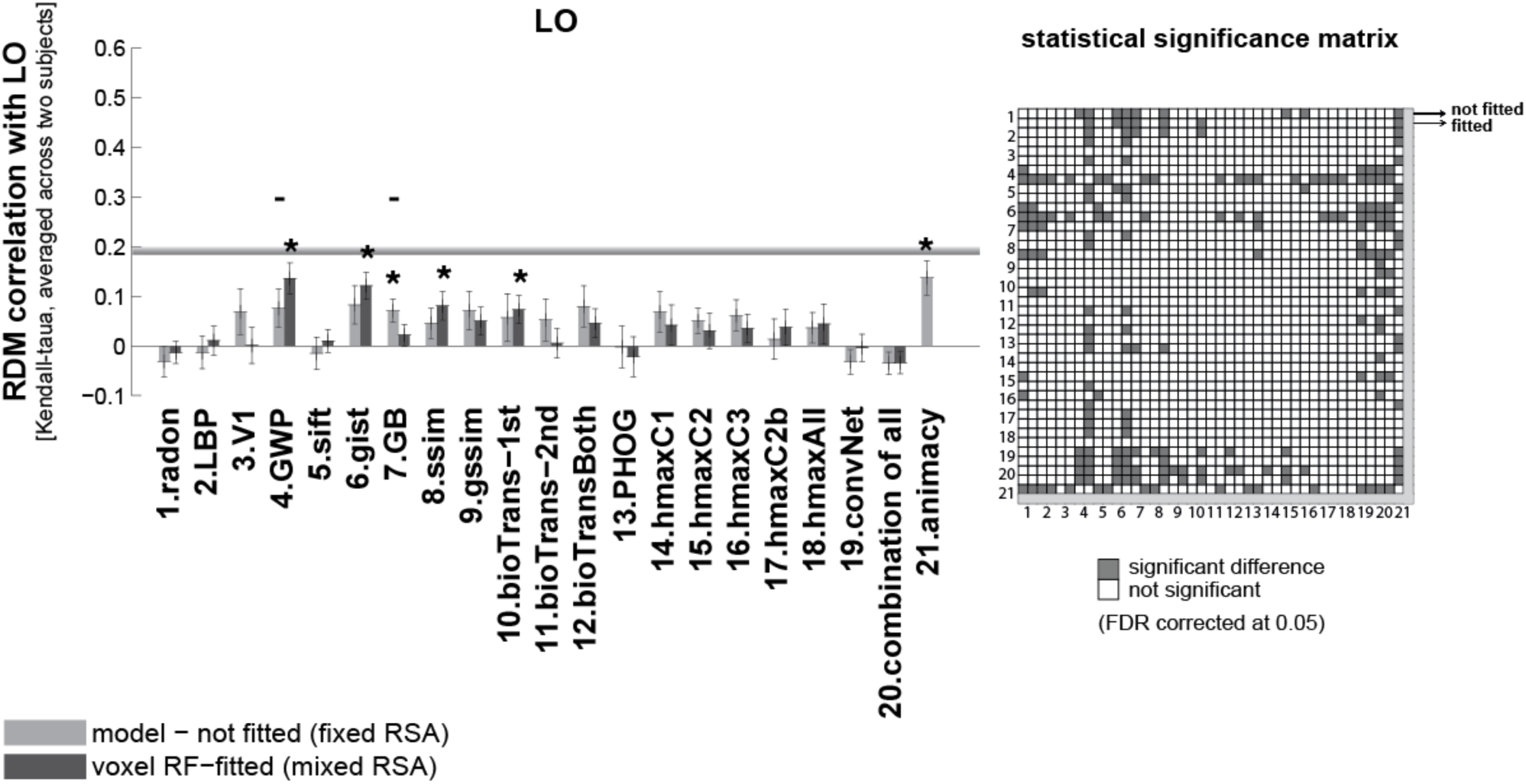
RDM correlation of unsupervised models and animacy model with higher visual area LO. Bars show the average of ten 12×12 RDM correlations (120 test stimuli in total) with the LO RDM. There are two bars for each model. The first bar, ‘model (not fitted)’, shows the RDM correlation of a model with a brain ROI without fitting the model responses to brain voxels (fixed RSA). The second bar (voxel RF-fitted) shows the RDM correlations of a model that is fitted to the voxels of the reference brain ROI, using 1750 training images (mixed RSA; refer to Figures 1 and 2 to see how the fitting is done). The gray horizontal line indicates the inter-subject RDM correlation in LO. The color-coded statistical signficance matrix at the right side of the bar graph shows whether any of the two models perform significantly differently in explaining LO (FDR corrected at 0.05). The statistical analyses and conventions here are analogous to Figure 4.

In 2012, a deep supervised convolutional neural network trained with 1.2 million labelled images (Krizhevsky et al., 2012) won the ImageNet competition (Deng et al., 2009) at 1000-category classification. It achieved top-1 and top-5 error rates on the ImageNet data that was significantly better than previous state-of-the-art results on this dataset. Following Khaligh-Razavi & Kriegeskorte (2014), we tested this deep supervised convolutional neural network, composed of 8 layers: 5 convolutional layers, followed by 3 fully connected layers. We compared the representational geometry of layers of this model with that of visual areas along the visual hierarchy (Figure 7).

**Figure 7.**
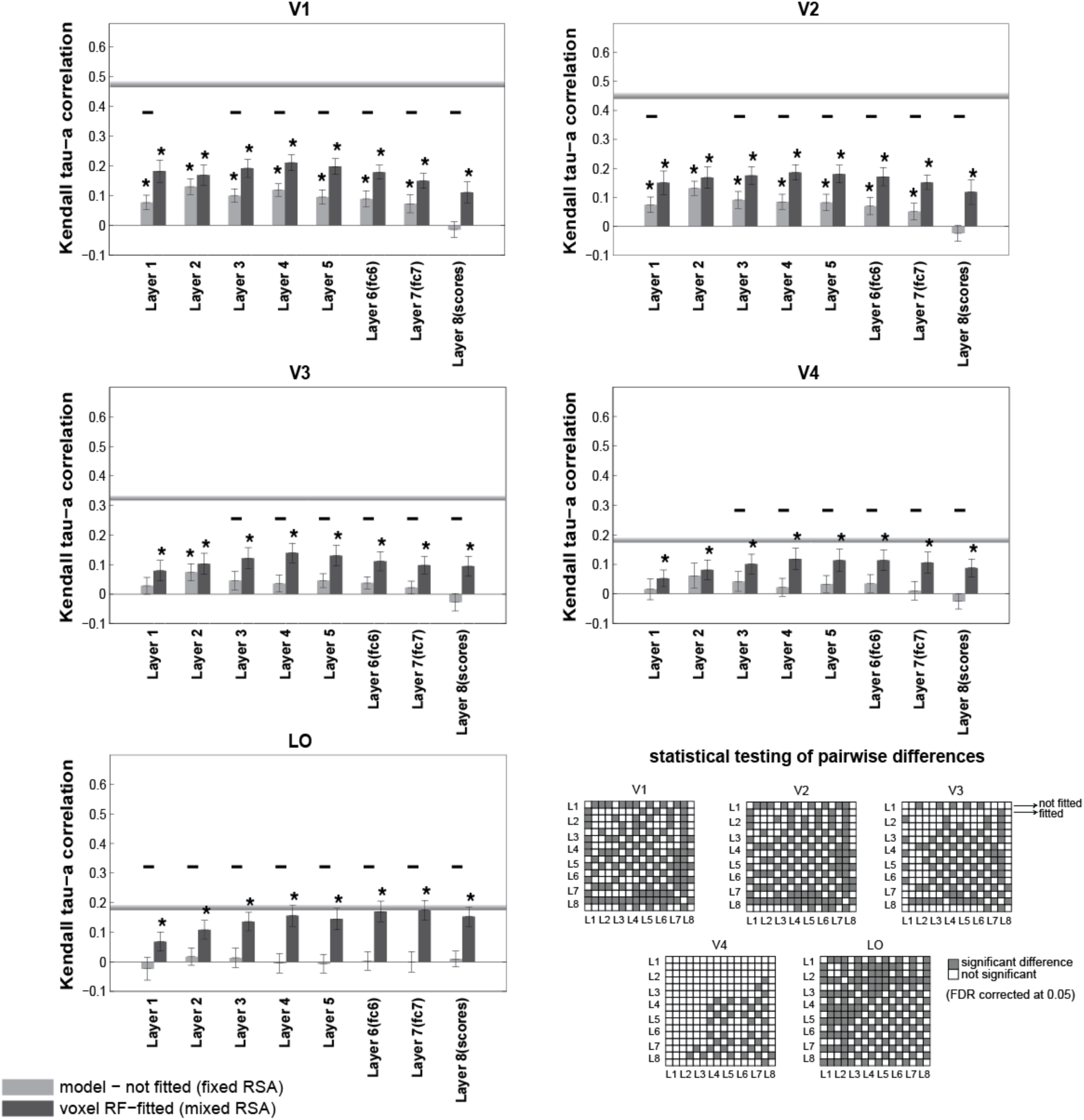
RDM correlation of the deep supervised convolutional network with brain ROIs across the visual hierarchy. Bars show the average of ten 12×12 RDM correlations (120 test stimuli in total) between different layers of the deep convolutional network with each of the brain ROIs. There are two bars for each layer of the model: the fixed RSA (model-not fitted), and the mixed RSA (voxel RF-fitted). The gray horizontal line in each panel indicates the inter-subject brain RDM correlation for the given ROI. The color-coded statistical signficance matrices show whether any of the two models perform significantly differently in explaining the corresponding reference brain ROI (FDR corrected at 0.05).The statistical analyses and conventions here are analogous to Figure 4.

Among all models, the ones that best explain LO are the mixed versions of layers 6 and 7 of the deep convolutional network. These layers also have a high animate/inanimate categorization accuracy (Figure 8B) – slightly higher than other layers of the network. Layer 6 of the deep net comes close to the inter-subject RDM correlation for LO as does the animacy model. The mixed version of some other layers of the deep convolutional network also come close to the LO inter-subject RDM correlation (Layers 3, 4, 5, and 8), as opposed to the unfitted versions. Remarkably, the mixed version of Layer 7 is the only model that reaches the inter-subject RDM correlation for LO.

**Figure 8.**
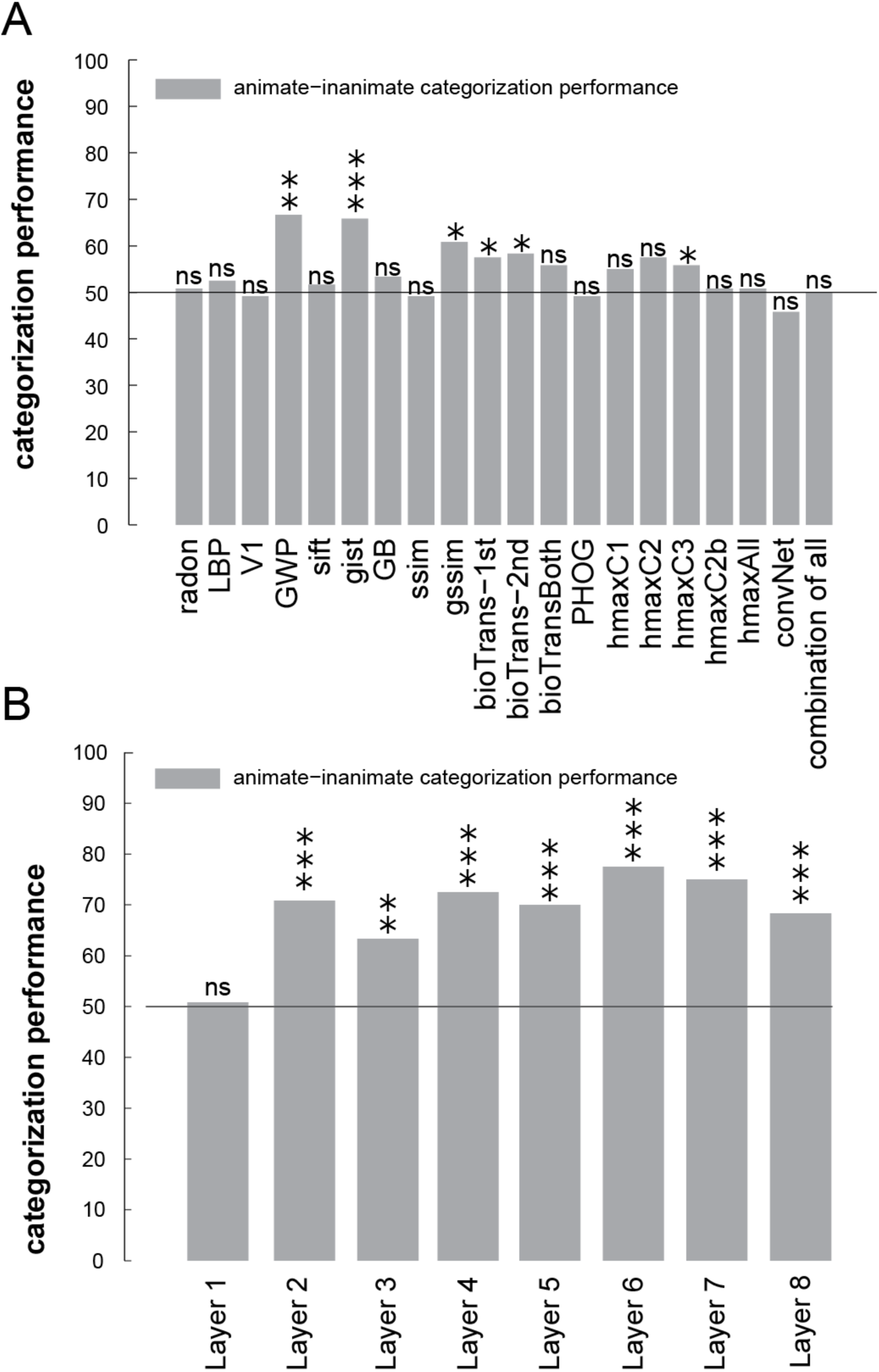
Animate-inanimate categorization performance for (A) several unsupervised models and (B) layers of a deep convolutional network. Bars show animate vs. inanimate categorization performance for each of the models shown on the X-axis. A linear SVM classifier was trained using 1750 training images and tested by 120 test images. P values that are shown by asterisks show whether the categorization performances significantly differ from chance [p < 0.05: *, p < 0.01: **, p < 0.001: ***]. P values were obtained by random permutation of the labels (number of permutations = 10,000).

Mixing brings consistent benefits to the deep supervised neural net representations (across layers and visual areas), but not to the shallow unsupervised models. For the deep supervised neural net layers (Figure 7), the mixed versions predict brain representations significantly better than the unmixed versions in 85% of the cases (34 of 40 inferential comparisons; 8 layers * 5 regions = 40 comparisons). For shallow unsupervised models (Figures 4–6), by contrast, the mixed versions predict significantly better in only 2% of the cases (2 of 100 comparisons; gist with V1, GWP with LO). For all other 98 comparisons (98%) between mixed and fixed unsupervised models, the fixed models either perform the same (e.g. ssim, LBP, SIFT) or significantly better than the mixed versions (e.g. HMAX, V1 model).

To asses the ability of object-vision models in the animate/inanimate categorization task, we trained a linear SVM classifier for each model using the model features extracted from 1750 training images (Figure 8). Animacy is strongly reflected in human and monkey higher ventral-stream areas (Kiani et al., 2007; Kriegeskorte et al., 2008a; Naselaris et al., 2012). We used the 120 test stimuli as the test set. To assess whether categorization accuracy on the test set was above chance level, we performed a permutation test, in which we retrained the SVMs on 10,000 (category-orthogonalized) random dichotomies among the stimuli. Light gray bars in Figure 8 show the model categorization accuracy on the 120 test stimuli. Categorization performance was significantly greater than chance for few of the unsupervised models, and all the layers of the deep ConvNet, except Layer1. Interestingly simple models, such as GWP and gist, also perform above chance at this task, though their performance is significantly lower than that of the higher layers of the deep network (Layers 6 and 7, p < 0.05).

Comparing the animate/inanimate categorization accuracy of the layers of the deep convolutional network (Figure 8B) with other models (Figure 8A) showed that the deep convolutional network is generally better at this task; particularly higher layers of the model perform better. In contrast to the unsupervised models, the deep convolutional network had been trained with many labelled images. Animacy is clearly represented in both LO and the deep net’s higher layers. Note, however, that the idealised animacy RDM did not reach the inter-subject RDM correlation for LO. Only the deep net’s Layer 7 (remixed) reached the inter-subject RDM correlation.

### Why does mixing help the supervised model features, but not the unsupervised model features?

Overall, fitting linear recombinations of the supervised deep net’s features gave significantly higher RDM correlations with brain ROIs than using the unfitted deep net representations (Figure 7). The opposite tended to hold for the unsupervised models (Figures 4–6). For example, the mixed features for all 8 layers of the supervised deep net have significantly higher RDM correlations with LO than the unmixed features (Figure 7). By contrast, only one of the unsupervised models (GWP) better explains LO when its features are mixed (Figure 6).

Why do features from the deep convolutional network require remixing, whereas the unsupervised features do not? One interpretation is that the unsupervised features provide general-purpose representations of natural images whose representational geometry is already somewhat similar to that of early and mid-level visual areas. Remixing is not required for these models (and associated with a moderate overfitting cost to generalization performance). The benefit of linear fitting of the representational space is therefore outweighed by the cost to prediction performance of overfitting. The deep net, by contrast, has features optimized to distinguish a set 1000 categories, whose frequencies are not matched to either the natural world or the prominence of their representation in visual cortex. For example, dog species were likely overrepresented in the training set. Although the resulting semantic features are related to those emphasized by the ventral visual stream, their relative prominence is incorrect in the model representation and fitting is essential.

This is consistent with our previous study (Khaligh-Razavi and Kriegeskorte, 2014), in which we showed that by remixing and reweighting features from the deep supervised convolutional network, we could fully explain the IT representational geometry for a different data set (that from Kriegeskorte et al. 2008). Note, however, that the method for mixing used in that study is different from the one in this manuscript as further discussed below in the the *Discussion* under ‘Pros and cons of fixed RSA, voxel-RF modeling, and mixed RSA’.

We know that a model (the voxel-receptive-field model here) might not generalize for a combination of two reasons:

1. Voxel-RF model parameters are overfitted to the training data. This is usually prevented or reduced by regularization. We did gradient descent with early stopping (which is a way of regularization) to prevent overfitting.
2. The model features do not span a representational space that can explain the brain representation. This is the problem of model misspecification. The model space does not include the true model, or even a good model.

In our case the lack of generalization does not happen in the deep net (in which we have many features), but it happens in some of the unsupervised models, which have fewer number of features than the deep net. (Figure 9 shows the number of features for each model.) The fact that fitting brings greater benefits to generalization performance for the models with more parameters is inconsistent with the overfitting account. Instead we suspect that the unsupervised models are missing essential nonlinear features needed to explain higher ventral-stream area LO.

**Figure 9.**
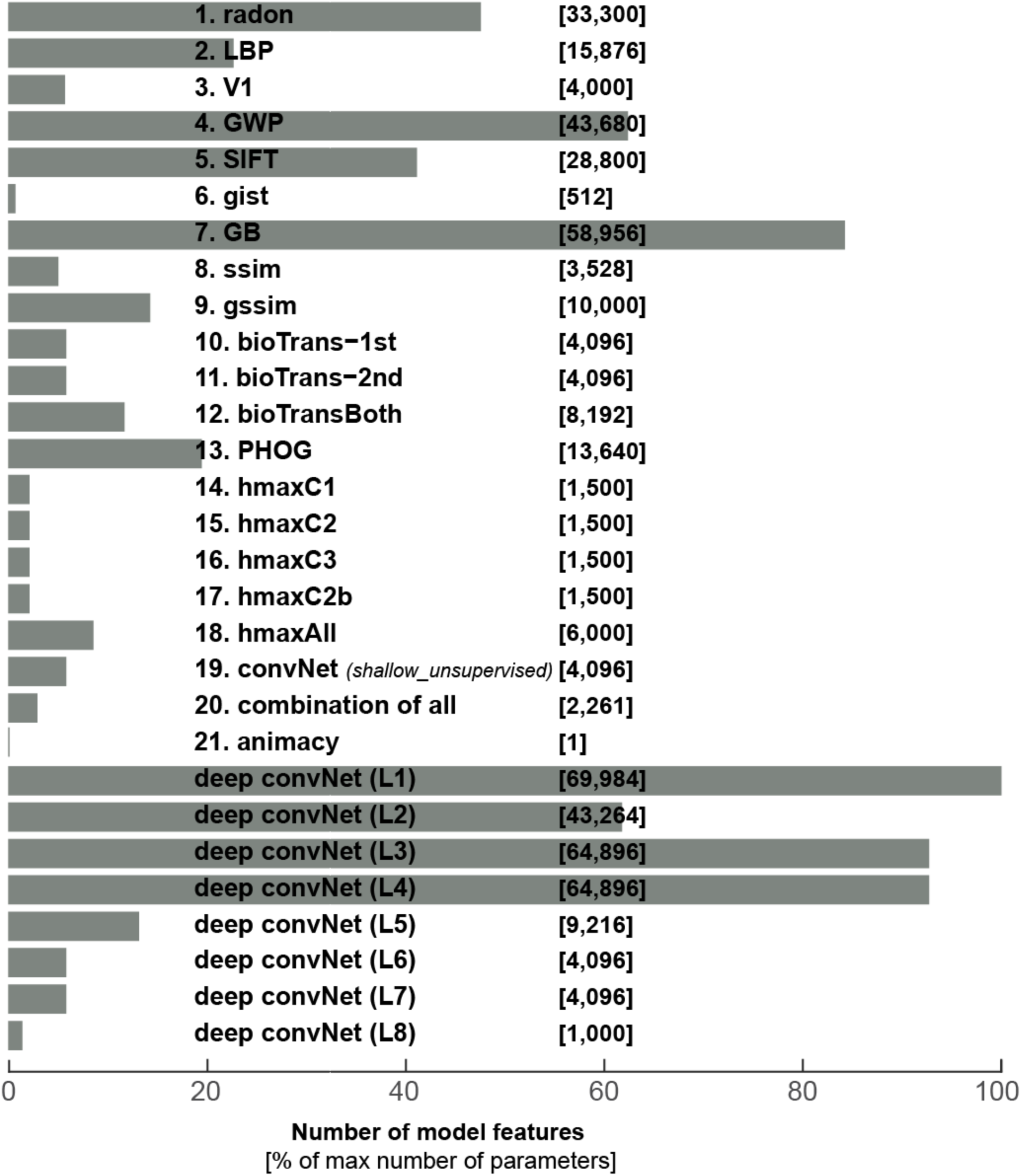
Number of features per model. The horizontal bars show percentage of max number of model features. The absolute number of model features are written in brackets in front of each model.The name for each model is written in front of its bar.

### Early layers of the deep convolutional network are inferior to GWP in explaining the early visual areas

Although the higher layers of the deep convolutional network successfully work as the best model in explaining higher visual areas, the early layers of the model are not as successful in explaining the early visual areas. The early visual areas (V1 and V2) are best explained by GWP model. The best layers of the deep convolutional network are ranked as the 4^th^ best model in explaining V1, and the 6^th^ best model in explaining V2. The RDM correlations of the first two layers of the deep convolutional network with V1 are 0.185 (Layer 1; voxel RF-fitted) and 0.18 (Layer 2; voxel RF-fitted), respectively. On the other hand, the RDM correlation of the GWP model (voxel RF-fitted) with V1 is 0.3, which is significantly higher than that of the early layers of the deep convNet (p < 0.001, signed-rank test). GWP appears to provide a better account of the early visual system than the early layers of the deep convolutional network. This suggests the possibility that improving the features in early layers of the deep convolutional network, in a way that makes them more similar to human early visual areas, might improve the performance of the model.

Güçlü and van Gerven, (2014) showed that shallow features learned from natural images with an unsupervised technique outperformed a Gabor wavelet model. In contrast to the present results, the model was trained without supervision, the architectures of the models and the method for comparing performance were different. In another study (Güçlü and Gerven, 2015), they found that a combination of CNN features (including features from lower layers and higher layers) explains early visual areas better than GWP. This is also different from our analysis here where we do not combine CNN features from different layers. Future studies should determine how early visual representations can be accounted for by shallow representations obtained by (1) predefinition, (2) unsupervised learning, and (3) supervised learning on various tasks within the same architecture.

### Mixed RSA and voxel receptive-field modelling yield highly consistent estimates of model predictive performance

We quantitatively compared the two methods for assessing model performances at explaining brain representations (Figure 10). The brain-to-model similarity was measured for all models using both mixed RSA and voxel-RF modeling. For mixed RSA, the brain-to-model similarity was defined as the Kendall tau-a correlation between model RDMs and brain RDMs on the 120 test stimuli. For voxel RF, the brain-to-model similarity was defined as the Kendall tau-a correlation between predicted voxel responses and the actual voxel responses. For each test stimulus, its predicted voxel responses are correlated with measured voxel responses, giving us 120 correlation values (for 120 test stimuli), which were then averaged. These values are shown for each model and each ROI in Figure 10 (values on the X-axis). In other words, in mixed RSA, the brain-to-model similarity is measured at the level of representational dissimilarities; and in voxel-RF model assessment, the brain-to-model similarity is measured at the level response patterns. We assessed the statistical significance of model-to-brain correlations using two-sided signed-rank test, corrected for multiple comparisons using FDR at 0.05. Models that did not significantly predict the representation in a given brain region fall within the transparent gray area (Figure 10).

**Figure 10.**
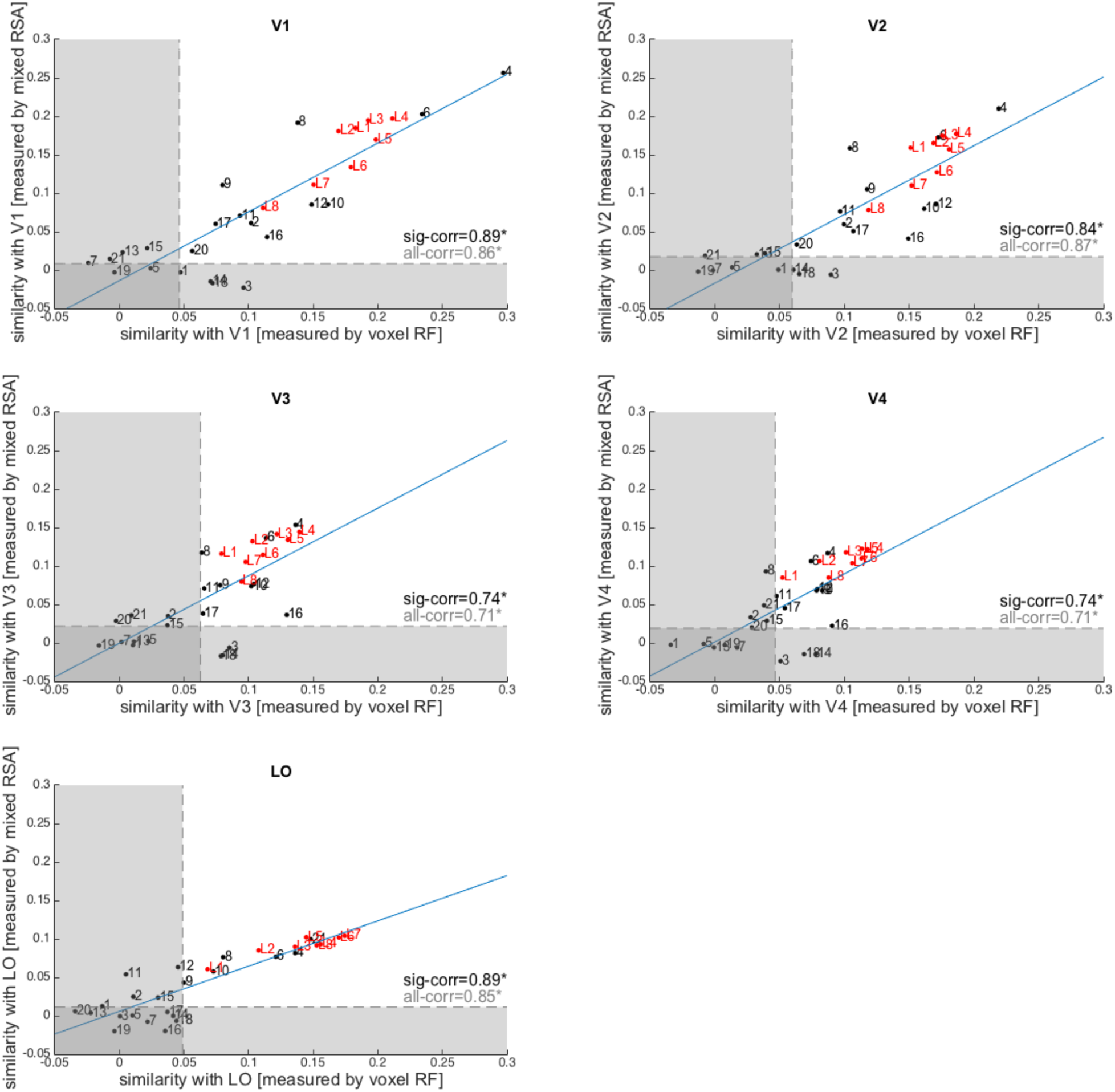
Mixed RSA and voxel-RF modelling are highly consistent. Each panel shows the consistency between mixed RSA and voxel RF in assessing model-to-brain similarity plotted for each ROI separately. Each dot is one model (red dots=deep CNN layers; black dots = unsupervised models). Numbers indicate the model (see Figure 9 for model numbering). The similarity measure with brain ROIs is Kendall tau-a correlation. For mixed RSA, it is the Tau-a correlation between the RDM of predicted voxel responses and a brain RDM (comparison at the level of dissimilarity patterns). For voxel RF, however, it is the Tau-a correlation between predicted voxel responses and actual voxel responses at the level of patterns themselves (averaged over 120 test stimuli). The transparent horizontal and vertical rectangles cover non-significant ranges along each axis (non-significant brain-to-model correlation). ‘sig-corr’ is the consistency between mixed RSA and voxel RF in assessing those models that fall within the significant range (outside the transparent rectangles). The consistency is measured as the Spearman rank correlation between mixed RSA brain-to-model correlations and voxel RF brain-to-model correlations. ‘all-corr’ is the consistency between mixed RSA and voxel RF across all models. Blue lines are the least square fits (using all dots).

The consistency between two approaches was defined as the Spearman rank correlation between the two sets of model-to-brain similarities. The model-to-brain similarities measured by these two methods were highly (Spearman r>0.7) and significantly correlated for all brain areas. In other words, the two approaches gave highly consistent results. Consistency was even higher for models explaining the brain representations significantly (Figure 10).

When a model fails to explain the brain data, comparing its mixed to its fixed performance with RSA provides a useful clue for interpretation of the model’s potential. Superior performance of the fixed model would indicate overfitting of the voxel-RF model. Failure of both mixed and fixed versions of a model suggests that the nonlinear features provided by the model are not appropriate (or that mixing is required, but the estimate is overfitted due to insufficient training data) (Figure 11).

**Figure 11.**
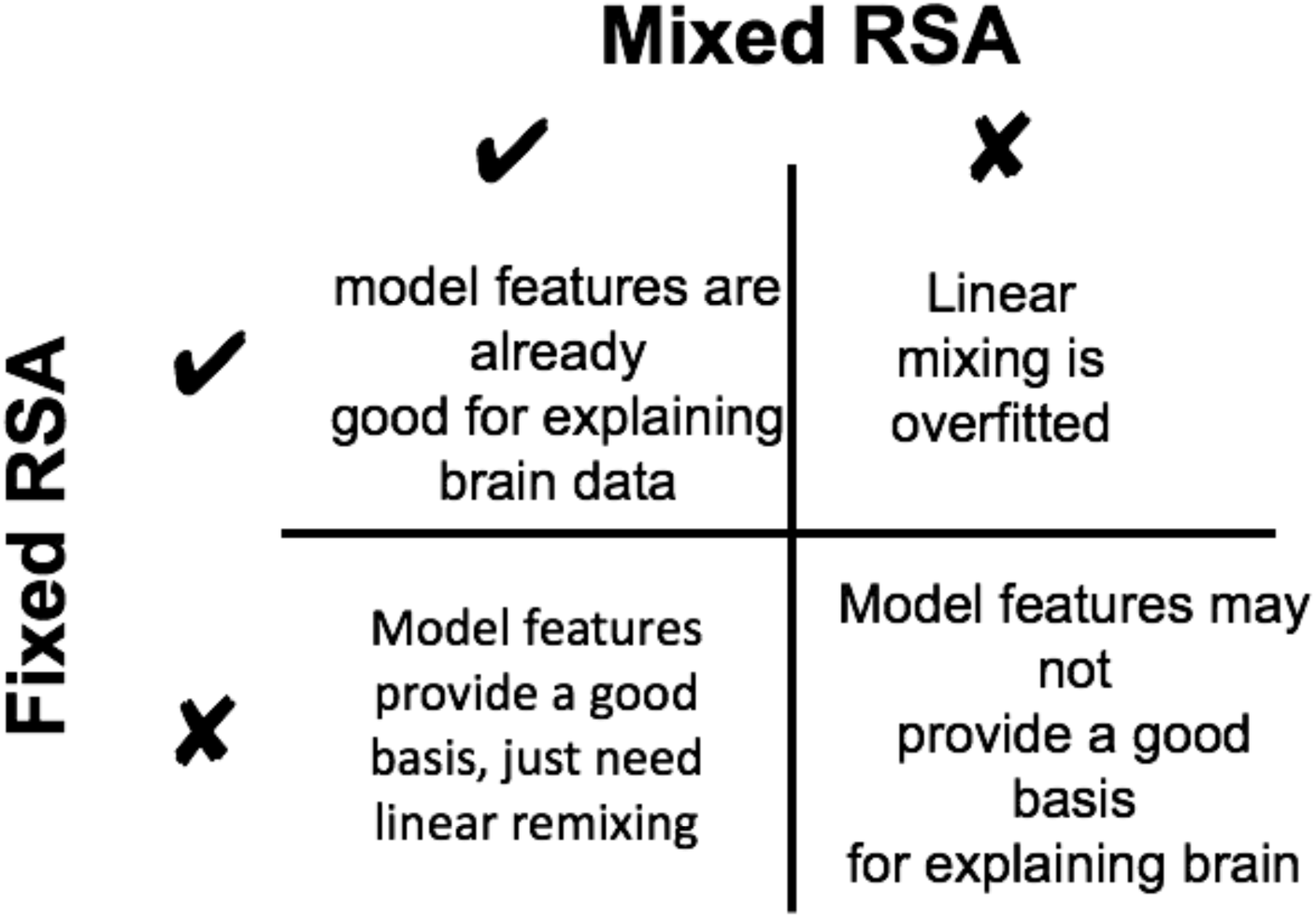
Mixed RSA vs. Fixed RSA. The figure shows how to interpret results from mixed RSA and fixed RSA. If a model explains brain data in both mixed and fixed RSA (top-left), this suggests that the model features are already good for explaining the brain data, obviating the need for feature remixing. If a model only explains brain data using fixed RSA but not the mixed RSA (top-right), it suggests that the linear mixing is overfitted. If on the other hand, a model only explains brain data using mixed RSA but not the fixed RSA (bottom-left), it shows that the model features provide a good basis for explaining the brain data and just need to be linearly remixed. Finally, if a model fails to explain brain data using both fixed and mixed RSA (bottom-right), the model features may not provide a good basis for explaining the brain data,

For example, HMAX-C2 (model number 15 in Figure 10) has an insignificant correlation with all brain ROIs in both mixed RSA and voxel-RF modeling. However, Figures 3 and 4 show that without mixing the fixed HMAX-C2 version explains brain representations significantly better. The raw HMAX-C2 features explain significant variance in the V1, V2, V3, and V4 representations. This would have gone unnoticed if analysis had relied only on the voxel-RF model predictions.

In summary, mixed RSA enables us to compare the predictive performance of the mixed and the fixed model. This provides useful information about (1) the predictive performance of the raw model representational space and (2) the benefits to predictive performance of fitting a linear mixing so as to best explain a brain representation (see Figure 11 for a summary).

## Discussion

Visual areas present a difficult challenge to computational modeling. We need to understand the transformation of representations across the stages of the visual hierarchy. Here we investigated the hierarchy of visual cortex by comparing the representational geometries of several visual areas (V1-4, LO) with a wide range of object-vision models, ranging from unsupervised to supervised, and from shallow to deep models. The shallow unsupervised models explained representations in early visual areas; and the deep supervised representations explained higher visual areas.

We presented a new method for testing models, mixed RSA, which bridges the gap between RSA and voxel-RF modelling. RSA and voxel-RF modelling have been used in separate studies for comparing computational models (Kriegeskorte et al., 2008a; Nili et al., 2014; Khaligh-Razavi and Kriegeskorte, 2014; Kay et al., 2008, 2013), but had not been either directly compared or integrated. Our direct comparison here suggests highly consistent results between mixed RSA and voxel-RF modeling – reflecting the fact that the same method is used to fit a linear transform using the training data. The difference lies in the level at which model predictions are compared to data: the level of brain responses in voxel-RF modeling and the level of representational dissimilarities in RSA. We also showed that a linearly mixed model can perform better (benefitting from fitting) or worse (suffering from overfitting) than the fixed original model representation. Comparing the predictive performance of mixed and fixed versions of each model provides useful constraints for interpretation.

### Pros and cons of fixed RSA, voxel-RF modeling, and mixed RSA

Voxel-RF modeling predicts brain responses to a set of stimuli as a linear combination of nonlinear features of the stimuli. One challenge with this approach is to avoid overfitting, through use of a sufficiently large training dataset in combination with prior assumptions that regularise the fit. This challenge scales with the number of model features, each of which requires a weight to be fitted for each of the response channels.

An alternative approach is RSA, which compares brain and model representations at the level of the dissimilarity structure of the response patterns (Kriegeskorte et al., 2008b; Kriegeskorte, 2009; Kriegeskorte and Kievit, 2013; Nili et al., 2014). This method enables us to test fixed models directly with data from a single stimulus set. Since the model is fixed, we need not worry about overfitting, and no training data is needed. However, if a model fails to explain a brain representation, it may still be the case that its features provide the necessary basis, but require linear remixing.

In Khaligh-Razavi & Kriegeskorte (2014), we had brain data for a set of only 96 stimuli. We did not have a large separate training set of brain responses to different images for fitting a linear mix of the large numbers of features of the models we were testing. To overcome this problem, we fitted linear combinations of the features, so as to emphasize categorical divisions known to be prevalent in higher ventral-stream representations (animate/inanimate, face/non-face, body/non-body). This required category labels for a separate set of training images, but no additional brain-activity data. We then combined the linear category readouts with the model features by fitting only a small number of prevalence weights (one for each model layer and one for each of the 3 category readout dimensions) with nonnegative least squares (see also, Jozwik et al., 2015). The fitting of these few parameters did not require a large training set comprising many images. We could use the set of 96 images, avoiding the circularity of overfitting (Kriegeskorte et al., 2009) by crossvalidation within this set.

In the present analyses, we had enough training data to fit linear combinations of model features in order to best explain brain responses. We first fit a linear model to predict voxel responses with computational models, as in voxel-RF modelling, and then constructed RDMs from the predicted responses and compared them with the brain RDMs, for a separate set of test stimuli. This approach enables us to test both mixed and fixed models, combining the strengths of both approaches.

Mixing enables us to investigate whether a linear combination of model features can provide a better explanation of the brain representational geometry. This helps address the question of whether the model features (a) provide a good basis for explaining a brain region and just need to be appropriately linearly combined or (b) the model features do not provide a good basis of the brain representation (see Figure 11 for a comparison between mixed and fixed RSA in this regard). However, mixing requires a combination of (1) substantial additional training data for a separate set of stimuli and (2) prior assumptions (e.g. implicit to the regularisation penalty) about the mixing weights. The former is costly and the latter affects the interpretation of the results, because the prior is part of the model.

The fact that mixed unsupervised models tended to perform worse than their fixed versions illustrates that mixing can entail overfitting and should not in general be interpreted as testing the best of all mixed models. If a large amount of data is available for training, mixed RSA combines the flexibility of the voxel-RF model fitting with the stability and additional interpretational constraint provided by testing fixed versions of the models with RSA.

### Performance of different models across the visual hierarchy

The models that we tested here were all feedforward models of vision, from shallow unsupervised feature extractors (e.g. SIFT) to a deep supervised convolutional neural network model that can perform object categorisation. We explored a wide range of models (two model instantiations for each of the 28 model representations + the animacy model = 57 model representations in total), extending previous findings (Khaligh-Razavi and Kriegeskorte, 2013, 2014) to the data set of Kay et al. (2008) and to multiple visual areas.

#### Fixed shallow unsupervised models explain the lower-level representations

The shallow unsupervised models explained substantial variance components of the early visual representations, although they did not reach the inter-subject RDM correlation. The mixed versions of the unsupervised models sometimes performed significantly worse than the original versions of those models (e.g. HMAX-C2). For lower visual areas, the fixed shallow unsupervised models appear to already approximate the representational spaces quite well.

For higher visual areas, the unsupervised models were not successful either with or without mixing. None of the unsupervised models came close to the LO inter-subject RDM correlation. One explanation for this is that these models are missing the visuo-semantic nonlinear features needed to explain these representations.

#### Mixed deep supervised neural network explains the higher-level representation

The lower layers of the deep supervised network performed slightly worse than the best unsupervised model at explaining the early visual representations. However, its higher layers performed best at explaining the higher-level LO representation. Importantly, the only model to reach the inter-subject RDM correlation for LO was the mixed version of Layer 7 of the deep net. Whereas the mixed versions of the unsupervised models performed similar or worse than the fixed versions, the mixed versions of the layers of the deep supervised net performed significantly better than their fixed counterparts.

A deep architecture trained to emphasize the right categorical divisions appears to be essential for explaining the computations underlying for the visuo-semantic representations in higher ventral-stream visual areas.

## Acknowledgements

We would like to thank Katherine Storrs for helpful comments on the manuscript. We would also like to thank all those who shared their model implementations with us. In particular Pavel Sountsov and John Lisman, who kindly helped us to set up their code. This work was supported by the UK Medical Research Council, a Cambridge Overseas Trust and Yousef Jameel Scholarship to SK; an Aalto University Fellowship Grant and an Academy of Finland Postdoctoral Researcher Grant (278957) to LH, and a European Research Council Starting Grant (261352) to NK. The authors declare no competing financial interests.

## Supplementary Methods

### Stimuli and response measurements details

Photographs were presented for 1 s with a delay of 3 s between successive photographs. The size of the photographs was 20° **x** 20° (500 pixels **x** 500 pixels). Subjects viewed the photographs while fixating a central white square. MRI data were collected at the Brain Imaging Center at University of California, Berkeley using a 4 T INOVA MR scanner (Varian, Inc.) and a quadrature transmit/receive surface coil (Midwest RF, LLC). For functional data, a T2*-weighted, single-shot, slice-interleaved, gradient-echo EPI pulse sequence was used: matrix size 64**x**64, TR 1 s, TE 28 ms, flip angle 20°. For anatomical data, a T1-weighted gradient-echo multislice sequence was used: matrix size 256**x**256, TR 0.2 s, TE 5 ms, flip angle 40°. Functional BOLD data were recorded from occipital cortex at a spatial resolution of 2mm **x** 2mm **x** 2.5mm and a temporal resolution of 1 Hz.

In this study, the data were pre-processed using an updated protocol (in comparison with the protocol in Kay et al., 2008). The new protocol included slice-timing correction, motion correction, upsampling to (1.5 mm)^3^ resolution and improved co-registration between the functional data sets. The data were modelled with a variant of the general linear model including discrete cosine basis set for the hemodynamic response function (HRF) estimation. Each basis function was 0 for the time point that is coincident with the onset of the image. Then, the basis functions extended 16 time points after that. Type-II DCT basis functions were used in the updated protocol (the first 7 functions were used). The beta weights characterizing the amplitude of the BOLD response to each stimulus were transformed to Z scores. Our analysis was restricted to voxels with signal-to-noise ratio greater than 1.5 (median value observed across all images).

Similar to (Kay et al. 2008) data from two subjects were analysed (S1–S2). The regions-of-interest (i.e. V1, V2, V3, V4, and LO) were identified using a retinotopic mapping procedure. The data for retinotopic mapping was collected in separate scan sessions. See (Kay et al., 2008; Naselaris et al., 2009; Kay et al., 2011) for further experimental details.

### Significance testing

For assessing statistical significance of RDM correlations, we used two-sided Wilcoxon signed-rank test, which is a non-parametric statistical test (Hollander and Wolfe, 1999; Gibbons and Chakraborti, 2011). Non-parametric tests do not make assumptions about the distribution of the data. The signed-rank test was used for testing the statistical significance of the RDM correlations between models and brain ROIs. To correct for multiple comparisons, where applicable, we used the false discovery rate (FDR) (Benjamini and Hochberg, 1995; Simmons et al., 2011). FDR-controlling procedures control the expected proportion of incorrectly rejected null hypotheses (“false discoveries”) among all rejected null hypotheses.

### Object-vision models

We used a wide range of computational models (for a review, see Khaligh-Razavi, 2014) to explore many different ways for extracting visual features. We selected some of the well-known bio-inspired object recognition models as well as several models and feature extractors from computer vision. Furthermore, to search the model space more comprehensively, in addition to the model representation itself, we made another instantiation from each model representation by linearly mixing the model features (i.e. voxel-RF fitted models). Below is a description for all models used in this study.

#### Gabor wavelet pyramid

The Gabor wavelet pyramid (GWP) model is similar to the GWP model used in Kay et al. (2008). Each image was represented by a set of Gabor wavelets of six spatial frequencies, eight orientations and two phases (quadrature pair) at a regular grid of positions over the image. To control gain differences across wavelets at different spatial scales, the gain of each wavelet was scaled such that the response of that wavelet to an optimal full-contrast sinusoidal grating is equal to 1. The response of each quadrature pair of wavelets was combined to reflect the contrast energy of that wavelet pair. The outputs of all wavelet pairs were concatenated to have a representational vector for each image.

#### Gist

The spatial envelope or gist model aims to characterize the global similarity of natural scenes (Oliva and Torralba, 2001). The gist descriptor is obtained by dividing the input image into 16 bins, and applying oriented Gabor filters in 8 orientations over different scales in each bin, and finally calculating the average filter energy in each bin^1^.

#### Animate–inanimate distinction

The natural images were labelled as animate if they contained one or several humans or animals, bodies of humans or animals, or human or animal faces. In the animate–inanimate model-RDM, the dissimilarities are either 0 (identical responses) if both images are of the same category (animate or inanimate) or 1 (different responses) if one image is animate and the other is inanimate. Because the animacy model is essentially one-dimensional, mixing it will not change the representational space beyond a scaling factor. Therefore, we did not include the mixed version of the animacy model.

#### Radon

The Radon transform of an image is a matrix, in which each column corresponds to a set of integrals of the image intensities along parallel lines of a given angle. The Matlab function Radon was used to compute the Radon transform for each luminance image.

#### Unsupervised convolutional network

A hierarchical architecture of two stages of feature extraction, each of which is formed by random convolutional filters and subsampling layers (Jarrett et al., 2009). Convolutional layers scan the input image inside their receptive field. Receptive Fields (RFs) of convolutional layers get their input from various places on the input image, and RFs with identical weights make a unit. The outputs of each unit make a feature map. Convolutional layers are then followed by subsampling layers that perform a local averaging and subsampling, which make the feature maps invariant to small shifts (Bengio et al., 1995). The convolutional network which we used^2^ had two stages of unsupervised random filters, that is shown by RR in table 1 in (Jarrett et al., 2009). The obtained result for each image was then vectorized. The parameters were exactly the same as used in (Jarrett et al., 2009).

#### Deep supervised convolutional neural network

The deep supervised convolutional network by Krizhevsky et al. (Donahue et al., 2013; Krizhevsky et al., 2012) is trained with 1.2 million labelled images from ImageNet (1000 category labels), and has 8 layers: 5 convolutional layers, followed by 3 fully connected layers. The output of the last layer is a distribution over the 1000 class labels. This is the result of applying a 1000-way softmax on the output of the last fully connected layer. The model has 60 million parameters and 650,000 neurons. The parameters are learnt with stochastic gradient descent.^3^

#### Biological Transform (BT)

BT is a hierarchical transform based on local spatial frequency analysis of oriented segments. This transform has two stages, each of which has an edge detector followed by an interval detector (Sountsov et al., 2011). The edge detector consists of a bar edge filter and a box filter. For a given interval *I* and angle *θ,* the interval detector finds edges that have angle *θ* and are separated by an interval *I*. In the first stage, for any given *θ* and *I*, all pixels of the filtered image were summed and then normalized by the squared sum of the input. They were then rectified by the Heaviside function. The second stage was the same as the first stage, except that in the first stage *θ* was changing between 0-180 ° and *I* between 100–700 pixels and the input to the first stage had not a periodic boundary condition on the *θ* axis (repeating the right-hand side of the image to the left of the image and vice versa); but in the second stage the input, which is the output of the first stage, was given a periodic boundary condition on the *θ* axis, and *I* was changing between 15-85 pixels.

#### Geometric Blur (GB)

289 uniformly distributed points were selected on each image, then the Geometric Blur descriptors(Belongie et al., 2002; Berg et al., 2005; Zhang et al., 2006) were calculated by applying spatially varying blur around the feature points. We used GB features that were part of multiple kernels for image classification described in (Vedaldi et al., 2009)^4^. The blur parameters were set to α=0.5 and β=1; the number of descriptors was set to 300.

#### Dense SIFT

For each grayscale image, SIFT descriptors (Lowe, 2004) of 16×16 pixel patches were sampled uniformly on a regular grid. Then, all the descriptors were concatenated in a vector as the SIFT representation of that image. We used the dense SIFT descriptors that were used in (Lazebnik et al., 2006) for extracting visual words.

#### Pyramid Histogram of Gradients (PHOG)

The canny edge detector was applied on grayscale images, and then a spatial pyramid was created with four levels (Bosch et al., 2007). The histogram of orientation gradients was calculated for all bins in each level. All histograms were then concatenated to create PHOG representation of the input image. We used Matlab implementation that was freely available online^5^. Number of quantization bins was set to forty, number of pyramid levels to four and the angular range to 360°.

#### Local Self-Similarity descriptor (ssim)

This is a descriptor that is not directly based on the image appearance; instead, it is based on the correlation surface of local self-similarities. For computing local self-similarity features at a specific point on the image, say *p*, a local internal correlation surface can be created around *p* by correlating the image patch centred at *p* to its immediate neighbours (Chatfield et al., 2009; Shechtman and Irani, 2007). We used the code available for ssim features that were part of multiple kernels for image classification described in (Vedaldi et al., 2009)^6^. The ssim descriptors were computed uniformly at every five pixels in both X and Y directions.

#### Global Self-Similarity descriptor (gssim)

This descriptor is an extension of the local self-similarity descriptor mentioned above. Gssim uses self-similarity globally to capture the spatial arrangements of self-similarity and long range similarities within the entire image (Deselaers and Ferrari, 2010). We used gssim Matlab implementation available online^7^. Number of clusters for the patch prototype codebook was set to 400, with 20000 patches to be clustered. D1 and D2 for the self-similarity hypercube were both set to 10.

#### Local Binary Patterns (LBP)

Local binary patterns are usually used in texture categorization. The underlying idea of LBP is that a 2-dimensional surface can be described by two complementary measures: local spatial patterns and gray scale contrast. For a given pixel, LBP descriptor gives binary labels to surrounding pixels by thresholding the difference between the intensity value of the pixel in the center and the surrounding pixels (Ojala et al., 2001, 2002; Pietikäinen, 2010). We used LBP Matlab implementation freely available online^8^. Number of sampling points was fixed to eight.

#### V1 model

A population of simple and complex cells were modelled and were fed by the luminance images as inputs. Gabor filters of 4 different orientations (0°, 90°, −45°, and 45°) and 12 sizes (7–29 pixels) were used as simple cell receptive fields. Then, the receptive field of complex cells were modelled by performing the MAX operation on the neighboring simple cells with similar orientations. The outputs of all simple and complex cells were concatenated in a vector as the V1 representational pattern of each image.

#### HMAX

The HMAX model developed by Serre et al.(Serre et al., 2007) has a hierarchical architecture inspired by the well-known simple to complex cells model of Huble & Wiesel (Hubel and Wiesel, 1968; HUBEL and WIESEL, 1962). There have been several extensions to the HMAX model, improving its feature selection process (e.g. Ghodrati et al., 2012) or adding new processing layers to the model (Zabbah et al., 2014). The HMAX model that is used here adds three more layers –ends at S4- on the top of the complex cell outputs of the V1 model described above. The model has alternating S and C layers. S layers perform a Gaussian-like operation on their inputs, and C layers perform a max-like operation, which makes the output invariant to small shifts in scale and position. We used the freely available version of the HMAX model^9^. All simple and complex layers were included until the S4 layer.

We used the pre-trained version of the HMAX model (i.e. trained with large number of natural images). Other versions of the HMAX model used in (Ghodrati et al., 2012; Rajaei et al., 2012) do not perform significantly better than the original version in terms if the similarity of their internal representations with brain ROIs. Therefore, we have only included the original version in this study. These models, and some other variations of the HMAX model have been compared in previous studies (Rajaei et al., 2012; Ghodrati et al., 2014a, 2014b; Khaligh-Razavi and Kriegeskorte, 2013, 2014; Ramakrishnan et al., 2014).

#### Combination of all

This is the concatenation of features extracted by the 19 unsupervised model representations. Given an input stimulus, features from all of the above-mentioned models were extracted. Because the dimension for extracted features differs across models, we used principle component analysis (PCA) to reduce the dimension of all of them to a unique number. We used the first 119 PCs from each of the models and concatenated them along a vector (119 was the largest possible number of PCs that we were able to use, because we had 120 testing images; so the covariance matrix has only 119 non-zero eigenvalues).

## Footnotes

1 http://people.csail.mit.edu/torralba/code/spatialenvelope/

2 http://koray.kavukcuoglu.org/code.html

3 http://caffe.berkeleyvision.org/

4 http://www.robots.ox.ac.uk/~vgg/software/MKL/#download

5 http://www.robots.ox.ac.uk/~vgg/research/caltech/phog.html

6 http://www.robots.ox.ac.uk/~vgg/software/SelfSimilarity/

7 http://www.vision.ee.ethz.ch/~calvin/software.html

8 http://www.cse.oulu.fi/CMV/Downloads/LBPMatlab

9 http://cbcl.mit.edu/software-datasets/pnas07/index.html

